## Supplemental Figures for "The double-stranded DNA-binding proteins TEBP-1 and TEBP-2 form a telomeric complex with POT-1"

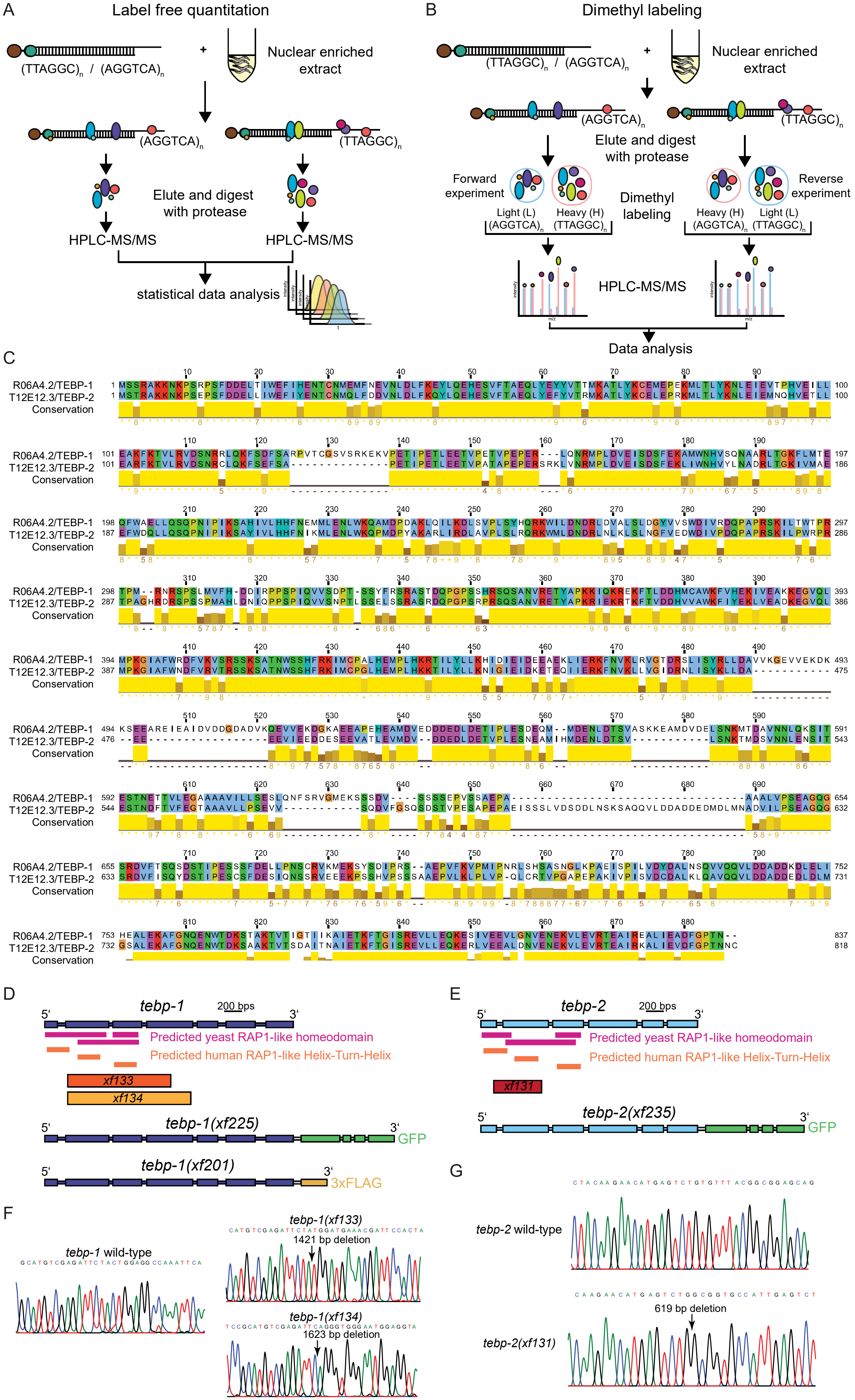

A
Label free quantitation
(TTAGGC)n / (AGGTCA)n
+
Nuclear enriched extract
(AGGTCA)n
(TTAGGC)n
Elute and digest with protease
HPLC-MS/MS
HPLC-MS/MS
statistical data analysis
B
Dimethyl labeling
(TTAGGC)n / (AGGTCA)n
+
Nuclear enriched extract
(AGGTCA)n
(TTAGGC)n
Elute and digest with protease
Forward experiment
Reverse experiment
Dimethyl labeling
Light (L) (AGGTCA)n
Heavy (H) (TTAGGC)n
Heavy (H) (AGGTCA)n
Light (L) (TTAGGC)n
HPLC-MS/MS
Data analysis
C
R06A4.2/TEBP-1
T12E12.3/TEBP-2
Conservation
D
tebp-1
200 bps
5'
3'
Predicted yeast RAP1-like homeodomain
Predicted human RAP1-like Helix-Turn-Helix
xf133
xf134
tebp-1(xf225)
GFP
tebp-1(xf201)
3xFLAG
E
tebp-2
200 bps
5'
3'
Predicted yeast RAP1-like homeodomain
Predicted human RAP1-like Helix-Turn-Helix
xf131
tebp-2(xf235)
GFP
F
tebp-1 wild-type
tebp-1(xf133)
1421 bp deletion
tebp-1(xf134)
1623 bp deletion
G
tebp-2 wild-type
tebp-2(xf131)
619 bp deletion


Figure S1

**Fig. S1. A quantitative proteomics screen to identify the paralogs TEBP-1 and TEBP-2.** (A) Scheme representing the label free quantitation workflow. Telomere $(TTAGGC)_n$, or control DNA $(AGGTCA)_n$ baits are incubated with nuclear extract. Samples are processed and measured independently, and later compared by statistical data analysis. (B) Scheme representing the reductive dimethyl labeling workflow. Telomere $(TTAGGC)_n$, or control DNA $(AGGTCA)_n$ baits are incubated with nuclear extract in duplicates. Per condition each peptide gets labeled with either light methyl groups ($CH_3$) or heavy methyl groups ($CD_3$). Afterwards, the heavy sample of one condition is combined with the light sample of the other condition and vice-versa to achieve a forward and a reverse experiment. Forward and reverse experiments are measured and analyzed by comparing intensities of the proteins (calculated from their peptide intensities) in the respective channel. (C) Pairwise sequence alignment of amino acid sequences of TEBP-1 and TEBP-2 using EMBOSS Needle, visualized using Jalview (2.11.0), showing the high sequence similarity between the two proteins. Amino acids are color coded according to the Clustal X colour scheme and conservation is shown in the yellow bars beneath the sequences. Amino acid positions are indicated. (D) Scheme of the *tebp-1* genomic locus. Below are indicated the positions with similarity to the homeodomain of human and yeast RAP1, as predicted by HHPred (3.2.0), deletions made by CRISPR-Cas9 genome editing (alleles *xf133* and *xf134*), as well as the locations of the tags (C-terminal GFP and 3xFLAG), also inserted by CRISPR-Cas9 genome editing. (E) As in (D) but for the *tebp-2* locus. (F-G) Chromatograms of Sanger sequencing of *tebp-1* and *tebp-2* deletion alleles compared to WT. Deletion sites are indicated with arrows.

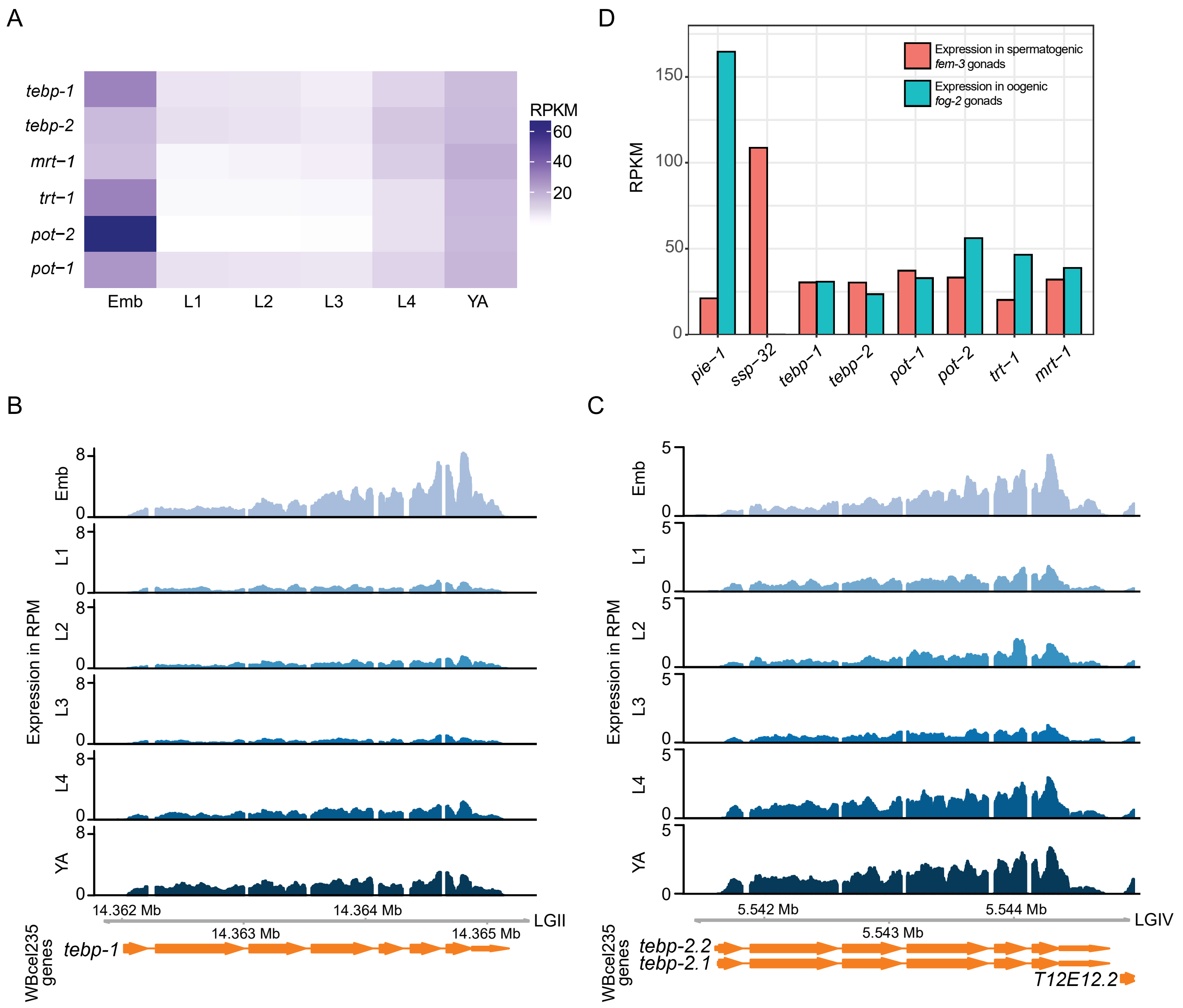
A
tebp-1
tebp-2
mrt−1
trt−1
pot−2
pot−1
Emb
L1
L2
L3
L4
YA
RPKM
60
40
20
D
RPKM
150
100
50
0
Expression in spermatogenic fem-3 gonads
Expression in oogenic fog-2 gonads
pie−1
ssp−32
tebp−1
tebp−2
pot−1
pot−2
trt−1
mrt−1
B
Expression in RPM
Emb
L1
L2
L3
L4
YA
8
0
14.362 Mb
14.363 Mb
14.364 Mb
14.365 Mb
LGII
WBcel235 genes
tebp-1
C
Expression in RPM
Emb
L1
L2
L3
L4
YA
5
0
5.542 Mb
5.543 Mb
5.544 Mb
LGIV
WBcel235 genes
tebp-2.2
tebp-2.1
T12E12.2

Figure S2

**Fig. S2. The expression profiles of *tebp-1* and *tebp-2* throughout development and in isolated gonads.** (A) Heatmap depicting mRNA expression levels, in Reads Per Kilobase Million (RPKM), of the known telomere binders *pot-1*, *pot-2*, and *mrt-1*, telomerase subunit *trt-1*, as well as *tebp-1* and *tebp-2*. Data from a previously published RNA-seq dataset [47]. (B-C) Genome browser tracks with the mRNA expression of *tebp-1* (B), and *tebp-2* (C), in counts per million (CPM), across the different life stages of *C. elegans*. Data from [47]. (A-C) Emb, embryos; L1-L4, first to fourth larval stages; YA, young adults. (D) Expression of telomere factors in dissected *fem-3* mutant gonads (exclusively spermatogenic) and *fog-2* mutant gonads (exclusively oogenic), from previously published RNA-seq data [48]. *pie-1* and *ssp-32* are genes known to be expressed in oogenesis and in spermatogenesis, respectively, according to [48].

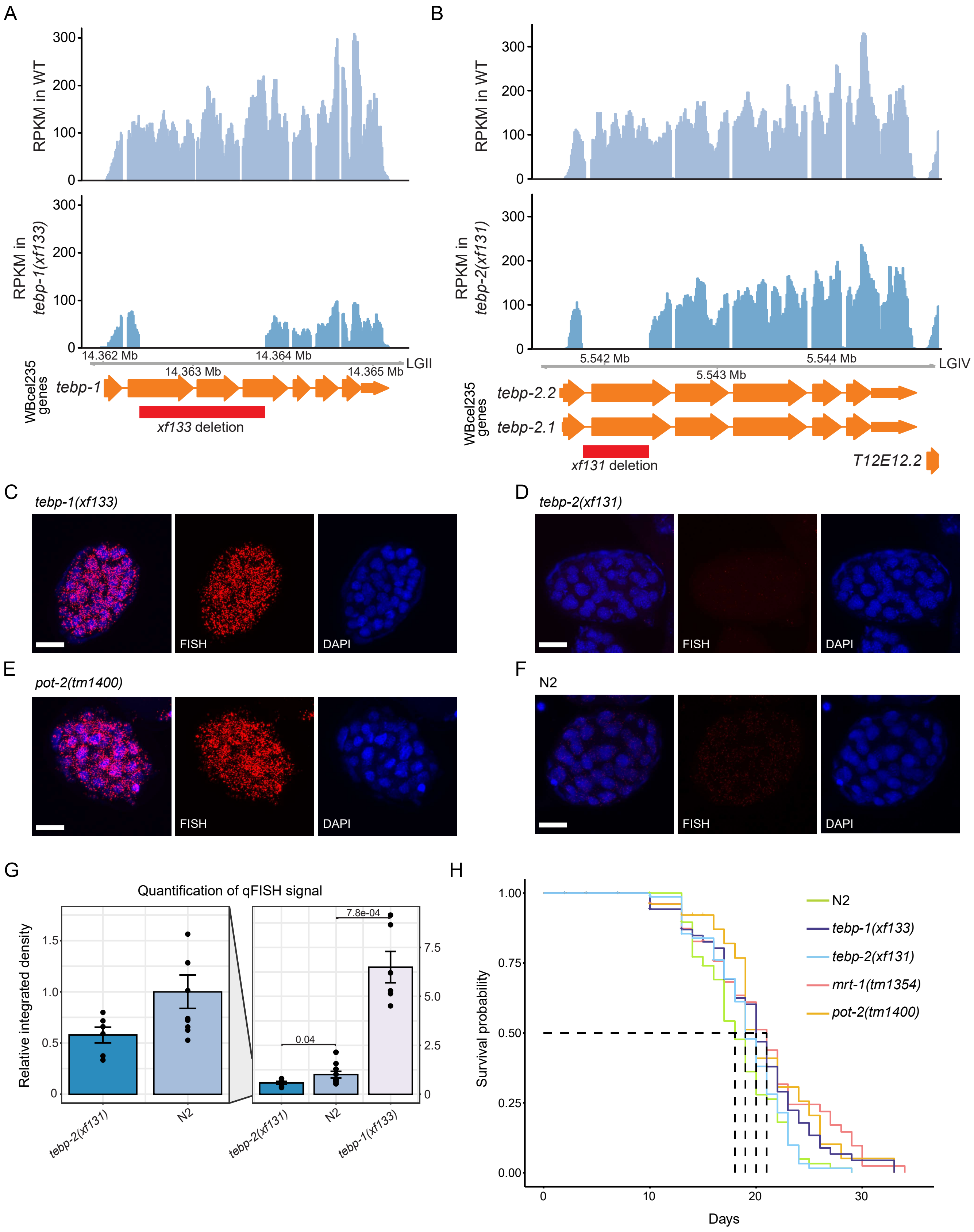
A
RPKM in WT
RPKM in tebp-1(xf133)
14.362 Mb
14.363 Mb
14.364 Mb
14.365 Mb
LGII
WBcel235 genes
tebp-1
xf133 deletion
B
RPKM in WT
RPKM in tebp-2(xf131)
5.542 Mb
5.543 Mb
5.544 Mb
LGIV
WBcel235 genes
tebp-2.2
tebp-2.1
xf131 deletion
T12E12.2
C
tebp-1(xf133)
FISH
DAPI
D
tebp-2(xf131)
FISH
DAPI
E
pot-2(tm1400)
FISH
DAPI
F
N2
FISH
DAPI
G
Quantification of qFISH signal
Relative integrated density
0.04
7.8e-04
tebp-2(xf131)
N2
tebp-1(xf133)
H
Survival probability
Days
N2
tebp-1(xf133)
tebp-2(xf131)
mrt-1(tm1354)
pot-2(tm1400)

Figure S3

**Fig. S3. TEBP-1 and TEBP-2 regulate telomere length in embryos but are dispensable for longevity.** (A-B) Genome browser tracks with the mRNA expression of *tebp-1* (A) and *tebp-2* (B), in Reads Per Kilobase Million (RPKM). RNA-seq data of wild-type, *tebp-1(xf133)*, and *tebp-2(xf131)* mutants. (C-F) Representative maximum projection z-stacks of a qFISH assay using embryos of *C. elegans* mutant strains. The telomeres of these embryos were visualized by hybridization with a telomeric PNA-FISH-probe. Nuclei were stained with DAPI. Scale bars, 10 µm. (G) Barplot depicting analysis of qFISH images of the strains in (C-F), as indicated on the x-axis. Average telomere length is indicated by arbitrary units of relative integrated density on the y-axis, with wild-type N2 set to 1. The left hand plot is a zoomed-in inset of the N2 and *tebp-2(xf131)* values. (H) Kaplan-Meyer plot showing the life span (in days) of animals of the indicated genotypes. L4 stage worms were picked for every strain and transferred every day, or every two days, to a fresh plate until death. n (excluding censored data, represented as crosses in the Fig.) as follows: N2, n=62; *tebp*-1, n=47; *tebp-2*, n=62; *pot-2*, n=21; *mrt-1*, n=42.

A

| Genotype | Father | Mother | Synthetic sterile F2 | Could grow double mutant homozygous line |
|---|---|---|---|---|
| *tebp-2(xf131); tebp-1(xf133)* | *tebp-2(xf131)* | *tebp-1(xf133)* | Yes | No |
| *tebp-1(xf133); tebp-2(xf131); xfIs148(tebp-2::gfp MosSCI)* | *tebp-1(xf133)* | *tebp-2(xf131); xfIs148(tebp-2::gfp MosSCI)* | No | Yes |
| *tebp-2(xf131); tebp-1(xf133)* | | | Yes | No |
| *tebp-2(xf131); pot-2(tm1400)* | *tebp-2(xf131)* | *pot-2(tm1400)* | No | Yes |
| *tebp-1(xf133); trt-1(ok410)* | *tebp-1(xf133)* | *trt-1(ok410)* | No | Yes |
| *tebp-1(xf133); mrt-1(tm1354)* | *tebp-1(xf133)* | *mrt-1(tm1354)* | No | Yes |
| *pot-2(tm1400); trt-1(ok410)* | *pot-2(tm1400)* | *trt-1(ok410)* | No | Yes |

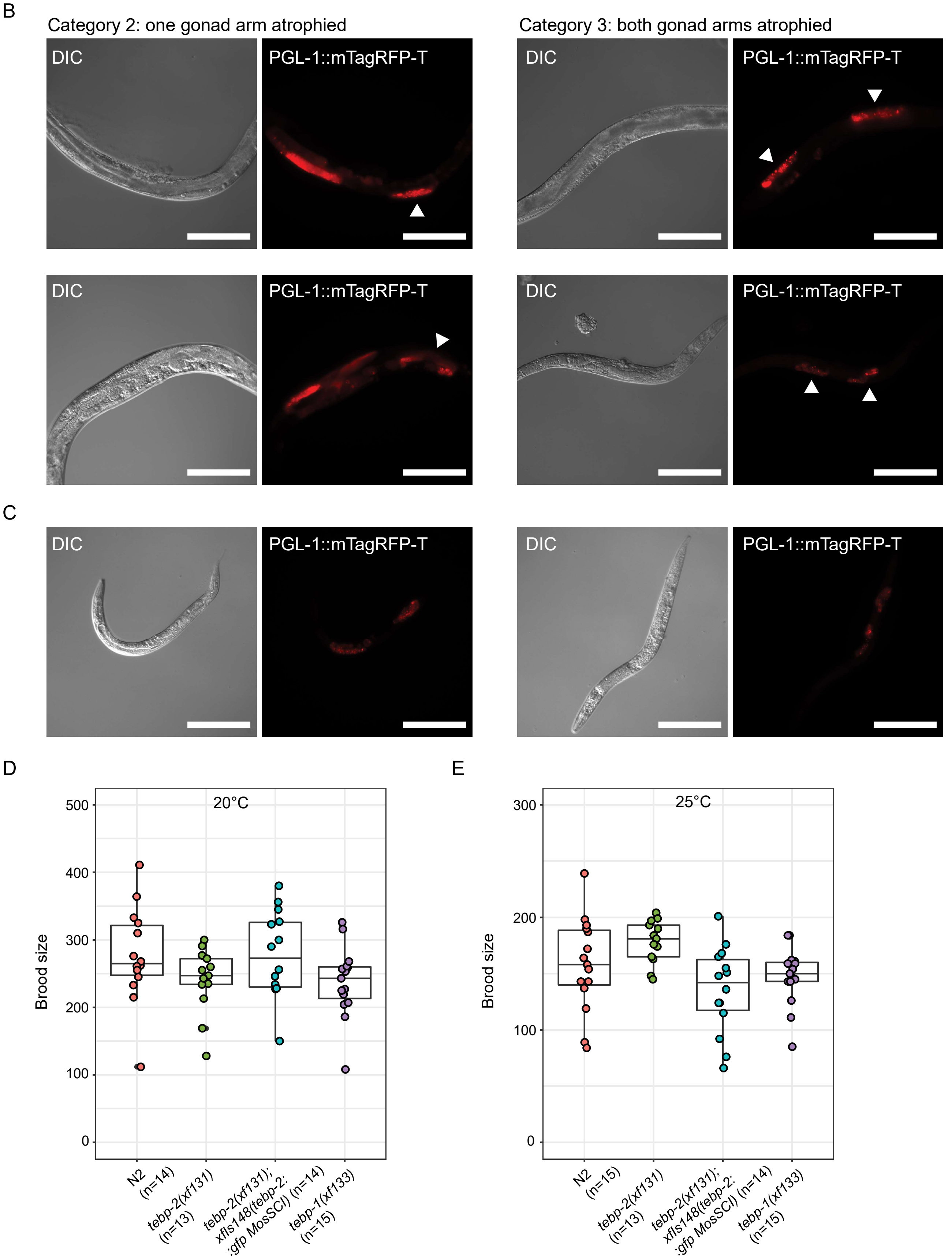


Figure S4

**Fig. S4. Dissecting the role of TEBP-1 and TEBP-2 in fertility. (A) Overview of additional crosses performed** to investigate distinct aspects of the synthetic sterility phenotype. For each cross, the columns indicate the genotype of the animals analyzed, the genotype of their parents, whether the animals have synthetic sterility, and if we could establish a homozygous line. The second row shows that the reciprocal cross between *tebp-1* and *tebp-2* also led to synthetic sterility. The third and fourth rows, show that a *tebp-2::gfp* single-copy transgene rescues the synthetic sterility of double mutants. The following rows demonstrate that the synthetic sterility is specific to *tebp-1* and *tebp-2*, as it does not arise in crosses with other telomere-associated mutants. (B) Additional representative widefield DIC and fluorescence pictures of worms with germlines of categories 2 (left panels) and 3 (right panels). Scale bars, 200 µm. Atrophied germlines are indicated with white arrowheads. (C) Exemplary widefield DIC and fluorescence micrographs of worms showing growth defects and/or larval arrest. These animals were isolated concurrently to animals shown in (B), but did not reach adulthood. These two specific animals were offspring of *tebp-2(xf131)*; *tebp-1(xf133)* +/-. Scale bars, 200 µm. (D-E) Boxplots showing the brood sizes of wild-type N2, *tebp-1* or *tebp-2* single mutants, and *tebp-2(xf131); xfIs148(tebp-2::gfp)*. Central horizontal lines represent the median, the bottom and top of the box represent the 25$^{th}$ and 75$^{th}$ percentile, respectively. Experiments were carried out at 20°C (D) and 25°C (E). Statistical comparisons were performed with wild-type N2, P-values>0.05, calculated with Mann–Whitney and Wilcoxon tests. n is indicated on the x-axis labels.

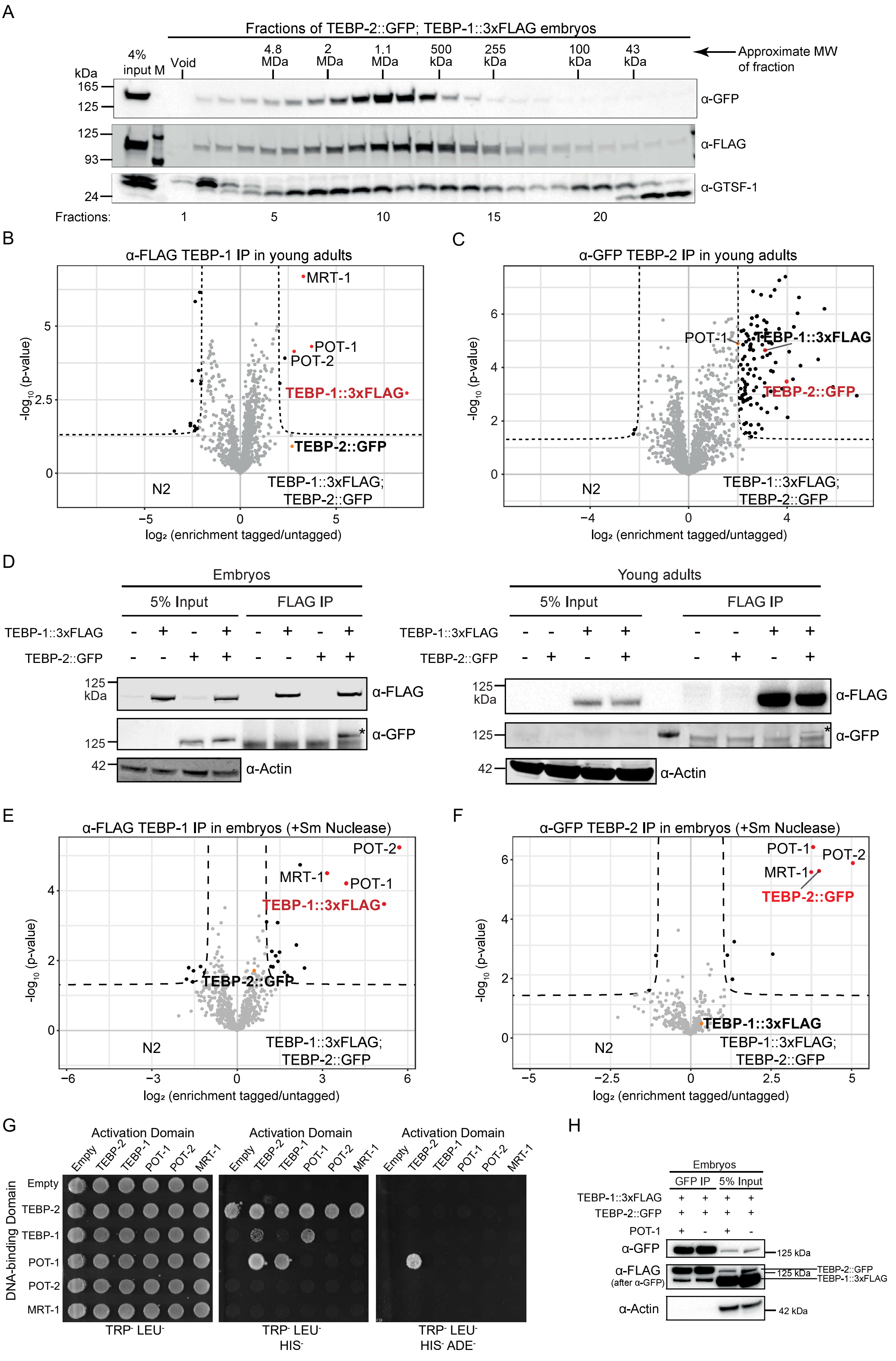
A
Fractions of TEBP-2::GFP; TEBP-1::3xFLAG embryos
Approximate MW of fraction
4% input
M
Void
4.8 MDa
2 MDa
1.1 MDa
500 kDa
255 kDa
100 kDa
43 kDa
kDa
165
125
93
24
α-GFP
α-FLAG
α-GTSF-1
Fractions:
1
5
10
15
20
B
α-FLAG TEBP-1 IP in young adults
MRT-1
POT-1
POT-2
TEBP-1::3xFLAG
TEBP-2::GFP
N2
TEBP-1::3xFLAG; TEBP-2::GFP
-log10 (p-value)
log2 (enrichment tagged/untagged)
C
α-GFP TEBP-2 IP in young adults
POT-1
TEBP-1::3xFLAG
TEBP-2::GFP
N2
TEBP-1::3xFLAG; TEBP-2::GFP
-log10 (p-value)
log2 (enrichment tagged/untagged)
D
Embryos
Young adults
5% Input
FLAG IP
TEBP-1::3xFLAG
TEBP-2::GFP
125 kDa
α-FLAG
α-GFP
α-Actin
42
E
α-FLAG TEBP-1 IP in embryos (+Sm Nuclease)
POT-2
MRT-1
POT-1
TEBP-1::3xFLAG
TEBP-2::GFP
N2
TEBP-1::3xFLAG; TEBP-2::GFP
-log10 (p-value)
log2 (enrichment tagged/untagged)
F
α-GFP TEBP-2 IP in embryos (+Sm Nuclease)
POT-1
POT-2
MRT-1
TEBP-2::GFP
TEBP-1::3xFLAG
N2
TEBP-1::3xFLAG; TEBP-2::GFP
-log10 (p-value)
log2 (enrichment tagged/untagged)
G
Activation Domain
Empty
TEBP-2
TEBP-1
POT-1
POT-2
MRT-1
DNA-binding Domain
TRP- LEU-
TRP- LEU- HIS-
TRP- LEU- HIS- ADE-
H
Embryos
GFP IP
5% Input
TEBP-1::3xFLAG
TEBP-2::GFP
POT-1
α-GFP
α-FLAG (after α-GFP)
α-Actin
125 kDa
TEBP-2::GFP
TEBP-1::3xFLAG
42 kDa

Figure S5

**Fig. S5. TEBP-1 and TEBP-2 interact with each other and with POT-1/MRT-1/POT-2.** (A) Western blot of the eluted fractions from size-exclusion chromatography of embryo extracts containing TEBP-1::3xFLAG and TEBP-2::GFP. The approximate molecular weight (MW) of the fractions is indicated above the blots. GTSF-1 was used as a control, as it has a known elution profile in size-exclusion chromatography [69]. Information about α-GTSF-1 can be found in [69]. (B-C) Volcano plots with quantitative proteomic analysis of TEBP-1::3xFLAG (B) or TEBP-2::GFP (C) IPs in young adults. IPs were performed in quadruplicates. Enriched proteins (threshold: 4-fold, p-value<0.05) are shown as black dots, enriched proteins of interest are highlighted with red or orange dots, and the baits are named in red. (D) Co-IP western blot experiment of TEBP-1::3xFLAG and TEBP-2::GFP similar to Fig. 5E-F, except the IPs were performed with an α-FLAG antibody. Actin was used as loading control. IPs with embryo extracts in the left panel and with young adult extracts in the right panel. (E-F) Volcano plots showing quantitative proteomic analysis of either TEBP-1::3xFLAG (E) or TEBP-2::GFP (F) IPs in embryos. IPs were performed in quadruplicates and Sm nuclease was added to remove potential DNA-dependent interactions. Enriched proteins (threshold >2-fold, p-value<0.05) are shown as black dots. Enriched proteins of interest are highlighted with red or orange dots, and baits are named in red. (G) Orthogonal grid of the Y2H spotting containing fusion constructs of the Gal4 activating or DNA-binding domains with the full length sequence of telomere factors. Left panel shows growth control in non-restrictive medium. Protein-protein interactions allow for growth on $TRP^-$ $LEU^-$ $HIS^-$ medium (middle panel). TEBP-2 bound to the Gal4 DNA-binding domain is self-activating, precluding the determination of interactions. The strongest interactions are permissive of growth on the highly stringent $TRP^-$ $LEU^-$ $HIS^-$ $ADE^-$ medium (right panel). (H) Co-IP western blot experiments of TEBP-1::3xFLAG and TEBP-2::GFP in the presence and absence of POT-1, where absence of POT-1 refers to the *pot-1(tm1620)* mutation. The IPs were performed with an α-GFP antibody. Actin was used as loading control. IPs were performed with 800 µg of embryo extracts. Detection by ECL was performed sequentially, first for GFP and then for FLAG.

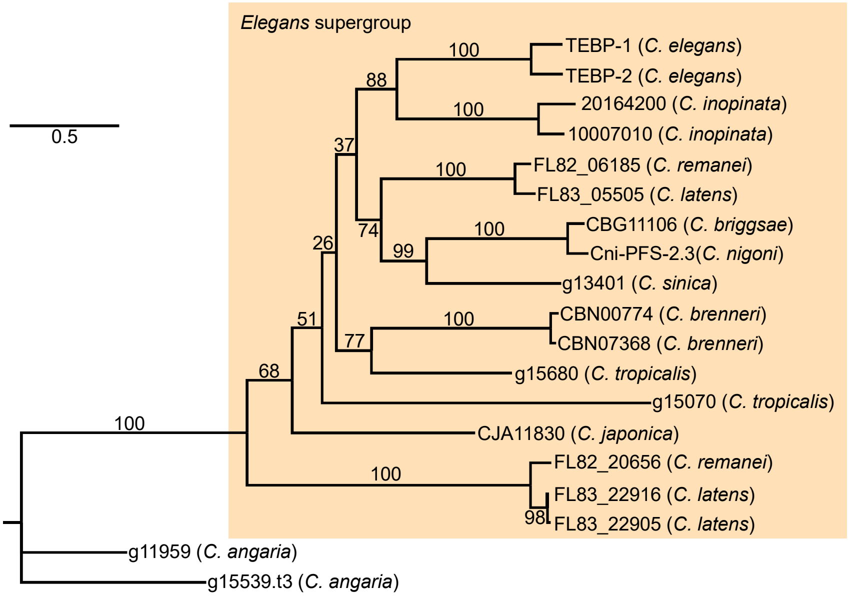
0.5
Elegans supergroup
100
TEBP-1 (C. elegans)
TEBP-2 (C. elegans)
88
100
20164200 (C. inopinata)
10007010 (C. inopinata)
37
100
FL82_06185 (C. remanei)
FL83_05505 (C. latens)
74
100
CBG11106 (C. briggsae)
Cni-PFS-2.3(C. nigoni)
26
99
g13401 (C. sinica)
100
CBN00774 (C. brenneri)
51
77
CBN07368 (C. brenneri)
g15680 (C. tropicalis)
68
g15070 (C. tropicalis)
100
CJA11830 (C. japonica)
100
FL82_20656 (C. remanei)
FL83_22916 (C. latens)
98
FL83_22905 (C. latens)
g11959 (C. angaria)
g15539.t3 (C. angaria)

Figure S6

**Fig. S6. Phylogenetic analysis of the N-terminal region of TEBP proteins.** Phylogenetic tree constructed as in Fig. 6A. The MAFFT protein alignment used for this tree comprised the first 600 alignment positions of the multiple sequence alignment in Supplementary Table 2 (sheet 2). Values on the nodes represent bootstrapping values of 10000 replicates, set to 100.
